# Human stem cell-based retina-on-chip as new translational model for validation of AAV retinal gene therapy vectors

**DOI:** 10.1101/2021.03.02.433550

**Authors:** Kevin Achberger, Madalena Cipriano, Matthias Düchs, Christian Schön, Stefan Michelfelder, Birgit Stierstorfer, Thorsten Lamla, Stefan G. Kauschke, Johanna Chuchuy, Julia Roosz, Lena Mesch, Virginia Cora, Selin Pars, Natalia Pashkovskaia, Serena Corti, Alexander Kleger, Sebastian Kreuz, Udo Maier, Stefan Liebau, Peter Loskill

## Abstract

Gene therapies using adeno-associated viruses (AAVs) are amongst the most promising strategies to treat or even cure hereditary and acquired retinal diseases. However, the development of new efficient AAV vectors is slow and costly, largely because of the lack of suitable non-clinical models. By faithfully recreating structure and function of human tissues, human induced pluripotent stem cell (iPSC)-derived retinal organoids could become an essential part of the test cascade addressing translational aspects. Organ-on-Chip (OoC) technology further provides the capability to recapitulate microphysiological tissue environments as well as a precise control over structural and temporal parameters. By employing our recently developed Retina-on-chip that merges organoid and OoC-technology, we analyzed the efficacy, kinetics and cell tropism of seven first and second generation AAV vectors. The presented data demonstrate the potential of iPSC-based OoC models as the next generation of screening platforms for future gene therapeutic studies.

## Introduction

Retinal diseases are the most common cause of visual impairment in developed countries and have become the worldwide leading cause of childhood blindness (Gilbert and Foster, 2001). Beside significant economic costs, these conditions have enormous consequences on the quality of life for the affected patients. Especially the absence of effective disease altering therapies and therapeutics resulting in poor prognosis create a high emotional burden.

Therefore, renewed efforts to find treatment options are constantly being undertaken. Despite being in the early stages, molecular diagnosis and new treatment strategies such as optogenetics, cell transplants and gene therapy already showed first promising results and foster hope for future efficacious treatment options (Wood, et al., 2019). In 2017, the first gene therapy for Leber’s congenital amaurosis was approved by the FDA (Trapani and Auricchio, 2018). In addition, novel gene therapeutic approaches for acquired diseases affecting larger patient populations are continuously developed.

Considered an easily accessible, highly compartmentalized and immune-privileged organ, the eye is a particularly promising target for gene therapy. Adeno-associated virus (AAV) vectors have become the gold standard for gene therapy addressing eye diseases. This is largely due to their favorable safety profiles, superior transduction capacity and long-lived gene expression (Trapani and Auricchio, 2018). To date, the majority of clinical trials deliver AAV vector via intravitreal or subretinal application (Ochakovski et al., 2017). Subretinal applications lead to a localized and highly efficient transduction of photoreceptors and the retinal pigment epithelium and are therefore used in most clinical trials addressing genetic disorders affecting these cell types. However, this complex procedure comes with a risk for complications in already compromised retinas due to retinal detachment during the operation (Garafalo et al., 2020). Conversely, the intravitreal application which is routinely used for application of therapeutics has the potential for a much broader biodistribution of applied vector and is considered as relatively safe. However, intravitreal injection with naturally occurring serotypes show overall poor transduction efficacy for retinal cells (Lipinski et al., 2013) resulting in low expression of therapeutic proteins and hence limit the potential for successful gene therapies. Capsid-modified, next-generation AAV vectors generated by screening of complex capsid libraries (Büning and Srivastava, 2019; Michelfelder and Trepel, 2009) are a promising approach to overcome the aforementioned lack of efficacy. However, the translation of results generated in preclinical models to the patient is still a major challenge (Dalkara et al., 2013; Hordeaux et al., 2018). So far, the field almost exclusively relied on mice as preclinical screening models in order to identify novel AAV vectors. Two promising approaches to overcome the lack of translatability are i) switching to non-human primates (NHPs) closer resembling the human physiology as screening models (Byrne et al., 2020), and ii) utilizing human-relevant *in vitro* models based on human (induced pluripotent stem cell (iPSC)-derived) retinal cells.

Self-assembling stratified organ-like tissues termed organoids derived from iPSCs have brought a new level of complexity to *in vitro* studies. Particularly retinal organoids (RO) constitute a prime example of the vast possibilities offered by the organoid technology. They contain all known major retinal cell types including photoreceptor cells, bipolar cells, Müller glia and ganglion cells and possess an *in vivo*-like retinal layering (Zhong et al., 2014). Most importantly, ROs build functional synaptic connections and are photosensitive. Recent studies assessing human fetal retina tissues and ROs side by side demonstrated the organoid’s close resemblance of retinal development and maturation to an advanced embryonic stage (Cowan et al., 2020).

The advantage of accurate *in vitro* manipulation and the human origin renders ROs very suitable for assessing the efficacy, cell toxicity and cell tropism in non-clinical screening models of gene therapy. However, the cultivation in suspension in dish culture has also major limitations. The poor media-to-tissue ratio, lack of vasculature and the unavoidable media change do not allow to study long-term effects of a given drug or gene therapeutics.

Besides relying on developmental biology to generate microphysiological tissues, microfabrication engineering provides a more controlled approach. By integrating human tissues into tailored microfluidic platforms, Organ-on-Chip (OoC) technologies recapitulate physiological tissue structure and function as well as vasculature perfusion. In the past decade, a variety of OoCs mimicking different types of tissues, organ functions and pathologies have been introduced (Marx et al., 2020; Zhang et al., 2018). Due to their human-relevance, amenability for experimental studies and high-content characteristic, they provide an immense potential for future drug development (Franzen et al., 2019; Low et al., 2020), especially in the field of ophthalmology (Haderspeck et al., 2019).

Still, one of the current limitations of OoC technology is the difficulty to generate complex stratified 3D tissues featuring a number of different cell types. This challenge can be addressed by incorporating organoids into OoC platforms. The integration of organoids into a controlled micro-environment of OoCs featuring vasculature-like perfusion and *in vivo* transport processes also solves major challenges of the organoid technology. By addressing the limitations of both technologies, the synergistic combination of OoCs and organoids paves the way for the next generation of stem cell-based microphysiological systems (Achberger et al., 2019; Takebe et al., 2017).

Here, we demonstrate for the first time that complex human iPSC-based OoCs can be utilized to test transduction efficacy of gene therapy in a pharmaceutical setting. Thereto, we employed a human Retina-on-Chip (RoC) model integrating iPSC-ROs and retinal pigmented epithelial cells (RPE) in a tailored microfluidic platform to test the performance of different types of AAV vectors. The compartmentalization and vasculature-like perfusion of the system enabled a physiological subretinal-like injection of the AAV particles and a nutrient supply via the choroidal-like vasculature. The optical accessibility provided the opportunity for *in situ* live-cell imaging. Additionally, we present results from initial screening steps on hiPSC-ROs providing higher throughput with lower complexity, a workflow which could serve as a blueprint for the drug development pipeline in future.

## Results

### AAV2, AAV2.7m8, and ShH10 show expression and cellular tropism after intravitreal application in mouse retina

First, performance of generated recombinant AAV vectors was analyzed in the eyes of adult mice. Therefore, different doses of AAV2 harboring either linear single-stranded (ssAAV2) or self-complementary (scAAV2) and AAV2.7m8 with self-complementary sequence of eGFP (scAAV2.7m8) under the control of the Cytomegalovirus (CMV) promotor were subjected to intravitreal application (**Fig 1**). 3 weeks after vector application, analyses of mRNA levels and histological stainings of the eye showed that all vectors expressed eGFP in a dose dependent manner (**Fig 1a**). Comparison of similar vector doses showed that switching from a linear single stranded eGFP sequence to a self-complementary sequence led to a marked increase in expression. Significantly higher eGFP levels were detected when sequences were delivered with the scAAV2.7m8 vs the scAAV2 capsid (**Fig 1a-b**). Of particular interest is the cell type-specific expression which can enable additional therapeutic approaches. ShH10 was described in rat models as preferentially transducing Müller glia cells after intravitreal application (Klimczak et al., 2009). In this study, histological analysis of mouse retinas for eGFP expression confirmed a strong cellular tropism of single-stranded ShH10 (ssShH10) vectors for Müller glia cells (**Fig 1c**). Transduction of Müller glia was observed applying a high (**Fig1c, i**) and low dose (**Fig1c, ii**) of ssShH10.

**Figure 1.**
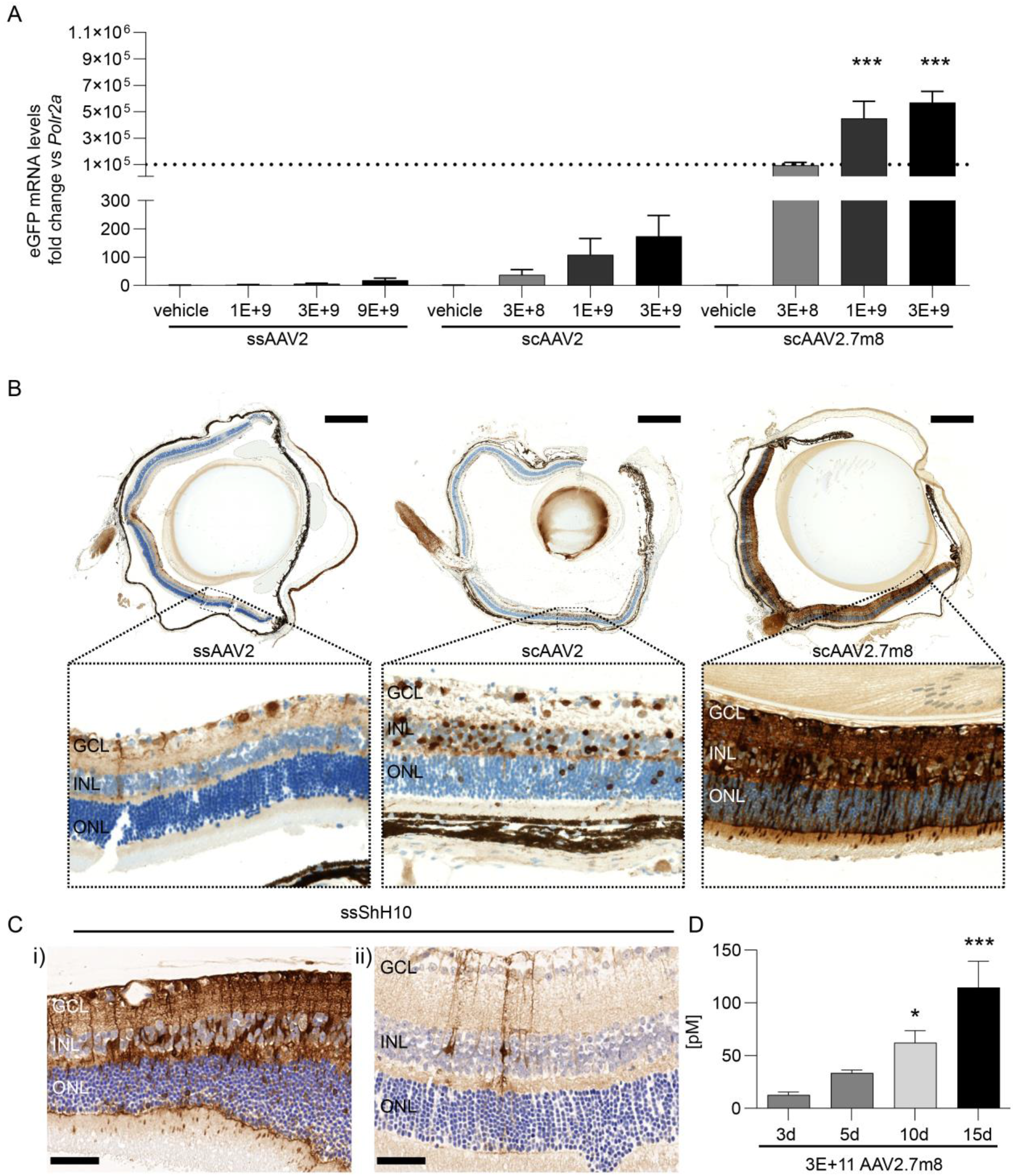
AAV induced expression in the mouse eye after intravitreal application.(A) eGFP RNA levels 3 weeks after injection. Statistical analysis in the graph represents the comparison with scAAV2 at similar doses. n=5-6 per condition, Mean ± SEM, ***p<0.001. (B) Representative vertical sections of mouse eyes three weeks after injection of 3E+9 vector genomes. Section were stained with anti-eGFP antibody (DAB, brown) and hematoxylin for cell nuclei (blue). Scale bars: (top) 500 µm, (bottom) 50µm. (C) Histological staining for eGFP in the mouse retina after i) 5E+9 vector genomes and ii) 1E+9 vector genomes of ssShH10. Scale bars: 50 µm. (D) Expression of secreted anti-FITC antibody over time course of 15 days. Statistical analysis represents comparison with day 3. n=5-6 per condition, Mean + SEM.

The constant increase of expression in the eye over the course of 15 days was verified by expression analysis of anti-FITC antibody protein in eye homogenate (**Fig 1d**). The sequence for the anti-FITC antibody was packed in a scAAV2.7m8 under the control of the CMV promotor Comparison of day 3 with day 10 and 15 showed a highly significant increase of anti-FITC antibody protein.

### AAV2, ShH10, AAV2.7m8 but not AAV9 efficiently transduce human ROs in vitro

As initial screening step, before moving into the RoC, we performed first line experiments in dish cultured ROs. For this proof of concept investigation, 4 AAV serotypes containing a self-complementary-CMV-eGFP expression cassette were selected: scAAV2, scAAV9, scShH10 and scAAV2.7m8.

As ROs show different cellular compositions depending on their age, experiments were performed in ROs with two defined developmental stages; day 80 ROs containing retinal progenitors, immature PRC, amacrine cells and ganglion cells, and day 300 ROs containing all retinal cell types with none or only rare observations of ganglion cells (Cowan et al., 2020).

ROs were imaged by brightfield and confocal high-resolution microscopy to assess general morphology and overall eGFP expression pattern (**Fig 2a**). The confocal setup provided a good estimate for expression profile at the surface of the ROs but is limited due their overall diameter (>500 µm). All quantifications of eGFP signals shown in this manuscript were done based on images acquired via standard fluorescence microscopy which allowed for detection of the overall fluorescence signal.

**Figure 2.**
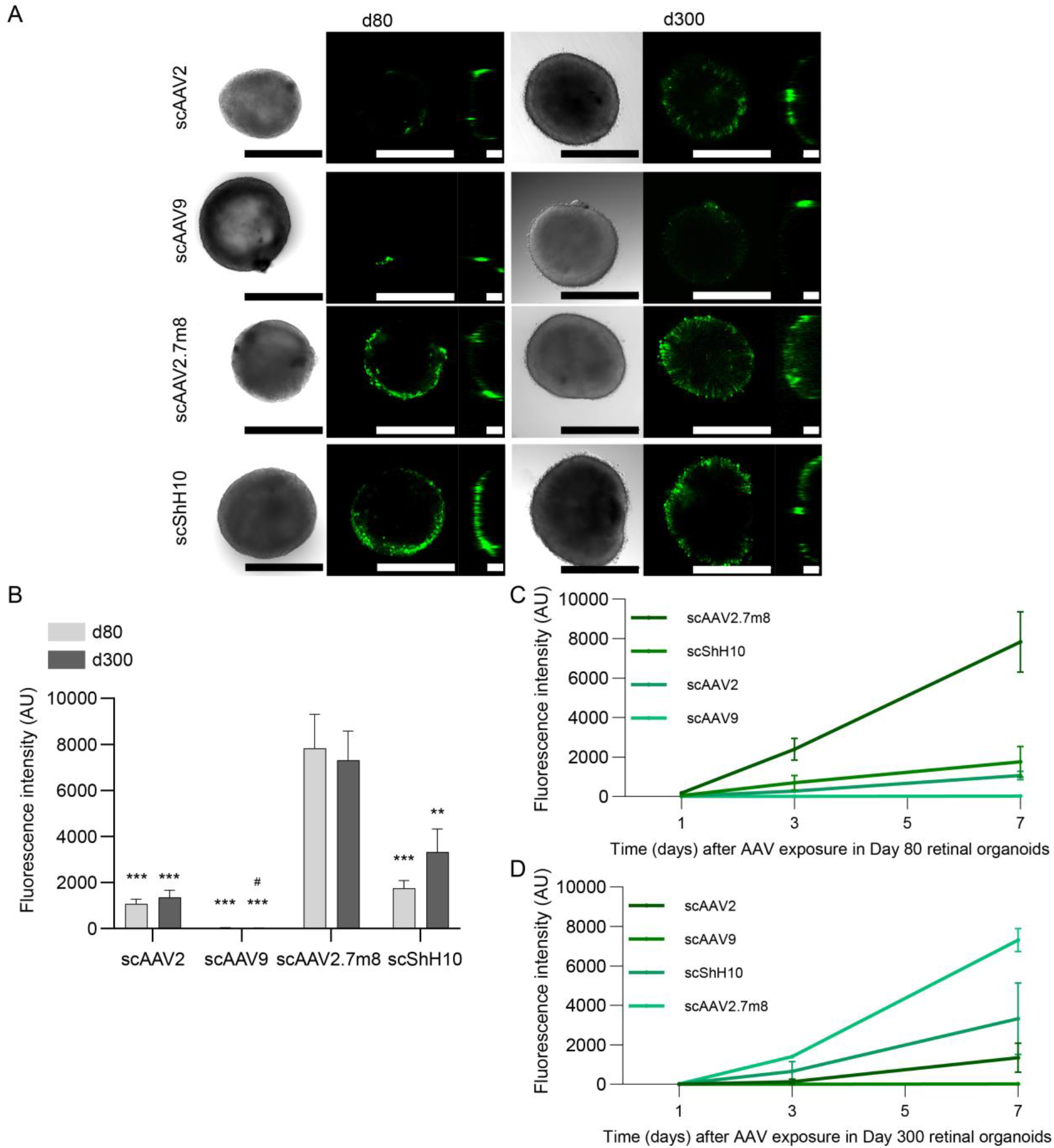
eGFP expression in AAV treated ROs differentiated for 80 and 300 days.(A) Confocal imaging of the AAV-treated organoids expressing eGFP (green). Left image: Brightfield, middle: xy-projection, right: yz-projection. Scale bars: 500µm. (B) Quantification of the mean eGFP fluorescence in day 80 and day 300 ROs exposed for 1, 2 or 3 days to the respective AAV (1E+10 virus genomes/well) and quantified after 7 days. Statistical analysis represents the comparison between AAV for the same RO age with scAAV2.7m8 (*) and with scShh10 (#). n= 17-18 per condition, Mean ± SEM. Kinetics of eGFP for (C) day 80 ROs and (D) day 300 ROs. n= 17-18 per condition, Mean ± SEM. See also **Figure S1-3**.

In the initial experiments, ROs were transduced in 48 well plates and incubation time with viral vector before media change was varied between 1 and 3 days. This, however, did not influence the eGFP signal measured 7 days after transduction (no statistical differences for all tested conditions; **Fig S1**). ScAAV2.7m8 showed the strongest overall eGFP expression after 7 days for both day 80 and day 300 ROs, when analyzing all incubation conditions side-by-side (1-3 days) (**Fig 2b & Fig S1**). A substantial fluorescence signal was also observed for scShH10. Here, day 300 ROs showed a slightly stronger signal than day 80 ROs. scAAV2 showed a comparably low expression for both day 80 and 300 ROs; scAAV9 signal was barely detectable in both conditions (**Fig 2b** and **Fig S1**). The comparison of signal magnitude at day 1, 3 and 7 post-transduction, showed distinct kinetics for each AAV serotype. With the exception of scAAV9, a time dependent increase of expression was observed for scAAV2, scShH10 and scAAV2.7m8. The steepest and overall strongest increase was induced by scAAV2.7m8 (**Fig 2c-d**).

Finally, we assessed if application of AAV vectors affects growth and morphology of the ROs (**Fig S2**). As expected, control day 300 ROs did not increase or decrease in size, since most of the cells have already become post-mitotic. In contrast to that, the size of the developmentally young RO of day 80 still increased around 20 % in controls and also in the low-transducing scAAV9. Interestingly, the size of ROs transduced with scAAV2, scShH10 and scAAV2.7m8 at day 80 decreased substantially 7 days after transduction suggesting either a loss of cells or overall RO integrity. This was supported by morphological assessment of brightfield images showing that most day 80 ROs transduced with scAAV2 (70 %), scShh10 (65 %) and scAAV2.7m8 (90 %) showed signs of degeneration or RO disorganization (**Fig 2a, Fig 2 S3a**). The observed morphological decline was proportional to the duration of viral incubation (**Fig S3b**). In contrast, day 300 ROs did not show any detrimental changes after viral transduction.

### The RoC as screening platform for evaluating AAV transduction efficiency

The RoC was recently developed to enable a stable co-culture and direct interaction of hiPSC-RPE and -RO while creating a tight outer blood-retina (epithelial) barrier (Achberger et al., 2019). The chip allows for a vascular like-perfusion through a bottom channel and a constant nutrient supply to the tissue-containing well through the epithelium (**Fig 3a**). This allows different compound application routes: systemic choroid-like application through the bottom channel and subretinal-like application into the wells. The latter allows to keep AAV particles present over a long period of time without nutrient deprivation mimicking a clinically practiced AAV application.

**Figure 3.**
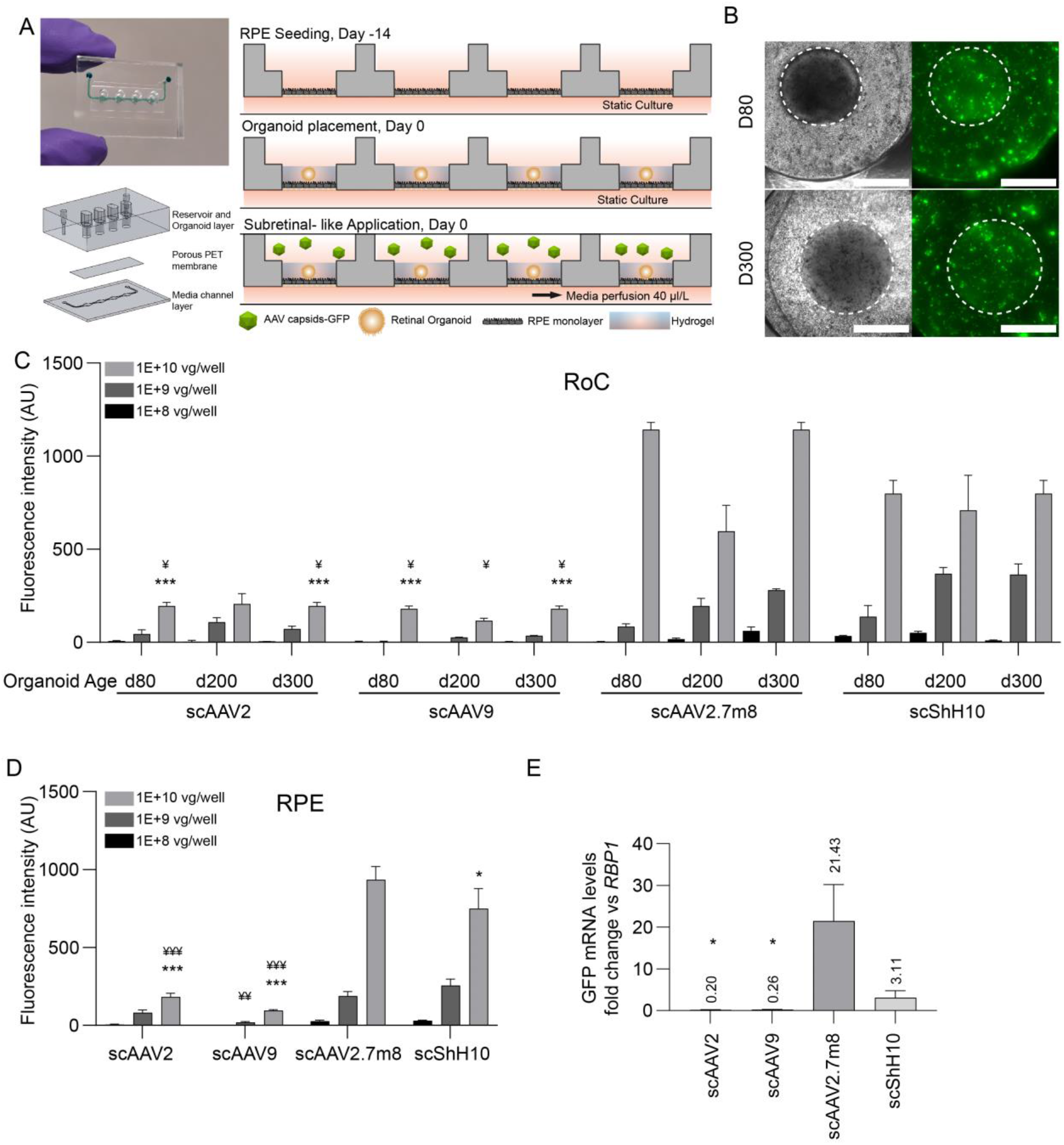
Testing of AAV serotypes in the RoC.(A) Adapted RoC design and chip seeding protocol for subretinal AAV-treatment. (B) Representative brightfield and GFP fluorescence live imaging. RoC area used for quantification is circled white. Fluorescent images are maximum intensity projections. Scale bars: 500µm. (C) Quantification of the mean eGFP fluorescence after 7 days of culture. Statistical analysis represents the comparison with scAAV2.7m8 (*) and scShH10 (¥) for 1E+10 virus genomes and the same organoid age. n=3 for all conditions, Mean + SEM. (D) Quantification of the mean eGFP fluorescence in the non-organoid areas of the RoC wells after 7 days of culture. Statistical analysis represents the comparison with scAAV9 (#), scAAV2.7m8 (*) and scShH10 (¥). n=9 per condition, Mean ± SEM, *p<0.05. (E) eGFP gene expression for each AAV in the RoC with day 300 ROs and a virus load of 1E+10 virus genomes per well. Statistical analysis represents the comparison with scAAV2.7m8. n=3-4 per condition, Mean ± SEM. See also **Figure S4 and S5**.

Each RoC contains four individual wells separated by a permeable membrane from the supply channel below (**Fig 3a**). To assemble the cellular components of the RoC, RPE cells are seeded in each well on top of the membrane on day -14. After 14 days of static cultures, ROs are seeded on top of the RPE and each well is infused with a hyaluronic acid-based hydrogel. Viral transduction occurred upon adding the AAVs to the well in a special 27µl compartment above the hydrogel. Thereafter, the chips were perfused with medium through the bottom channel while keeping the viral vector-containing upper well static.

For this study, ROs of three ages (days 80, 200 and 300) were selected and three different doses of viral vector (1E+10, 1E+9, 1E+8 vector genomes per RoC well) were applied. After 3 and 7 days, total eGFP fluorescence of RO area and the underlying RPE (refered as RoC area, circled white in **Fig 3b**) of each AAV serotype was assessed. 7 days post-transduction, a clear dose to signal correlation was confirmed for all conditions and AAV serotype (**Fig 3c**). ScAAV2.7m8 and scShH10 induced the strongest signals throughout the three different RO ages (**Fig 3c**), significantly outperforming scAAV2 and scAAV9. Both, scAAV2 and scAAV9, only induced a weak expression in all RoC irrespective of RO age. When assessing the AAV transduction in the non-organoid, solely RPE cell containing area, again scAAV2.7m8 and scShH10 led to significantly higher signals than scAAV2 and scAAV9 (**Fig 3d)**. In the following, spatial distribution of the eGFP signal was visualized using confocal imaging (**Fig S4**). We found that both RPE and RO contributed to the overall eGFP fluorescence, with the strongest eGFP signal found in the scAAV2.7m8-transduced RoC. This finding was supported by the assessment of eGFP mRNA levels of RoC-extracted ROs (**Fig 3e**) in which again scAAV2.7m8 by far induced the highest expression levels with a 21-fold increase (relative to RNA polymerase II) in ROs, followed by ShH10 (3.11), AAV9 (0.26) and AAV2 (0.20). Finally, day 7 to day 3 comparison was used to assess the AAV serotype-specific kinetics (**Fig S5**). These analyzes confirmed the ability to capture distinct transduction profiles of single AAV vectors in the RoC.

### AAV serotypes show wide range cell tropism in the RoC

Next, we interrogated cell tropism of different AAV serotypes within the human ROs. RoCs containing day 200 ROs were transduced with 1E+10 virus genomes, day 80 ROs were transduced with 1E+09 virus genomes. 7 days after AAV application, ROs were harvested, cryosectioned and co-stained with cell type specific markers such as GNAT1 (rod photoreceptors), ARR3 (cone photoreceptors), CRALBP (Müller glia) and BRN3B (ganglion cells) (**Fig 4**). Our investigations indicate that all 4 AAVs were associated with a tropism for rod and cone photoreceptors as well as Müller glia cells in day 200 ROs (**Fig 4a-c**). This cell tropism was recapitulated at day 300 ROs (data not shown). Ganglion cell tropism for all 4 serotypes could be shown in ROs at day 80 (**Fig 4d**). Furthermore, for amacrine, horizontal and bipolar cells, we collectively found positively co-stained cells for all serotypes with the only exception of scAAV2 which did not transduce amacrine cells in our setup (**Figure S6**).

**Figure 4.**
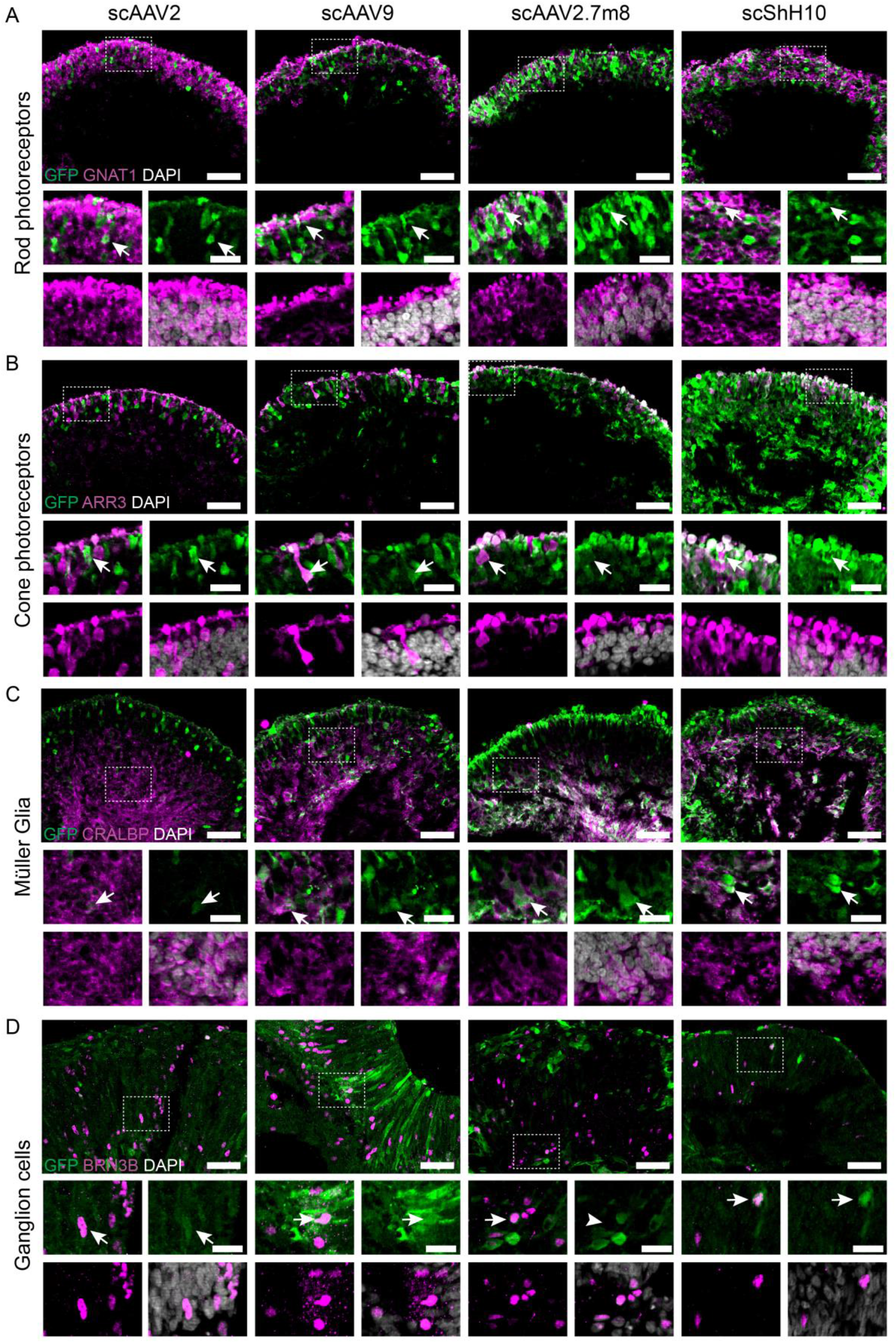
Evaluation of cell tropism in the RoC. Day 200 ROs in (A-C) at were transduced with 1E+10 virus genomes, (D) day 80 ROs were transduced with 1E+9 virus genomes. (A-D) Vertical cryo-sections of ROs showing AAV-mediated eGFP expression (green) and cellular co-stainings (magenta). Hoechst: white. Co-stained cells are highlighted with white arrow. Scale bars: 50 µm (large images), 20 µm (small images). See also **Figure S6**.

### Screening of next generation AAVs in the RoC

Aiming to optimize tropism and transduction efficiency, new AAVs are constantly generated. To show the applicability of the RoC as a screening system for novel AAV variants, we applied two recently developed 2^nd^ generation AAV capsids: AAV2.NN and AAV2.GL (Pavlou et al., 2021). Both variants show highly efficient transduction profiles for various retinal cell types (Pavlou et al., 2021). Here, we analyzed scAAV2.NN and scAAV2.GL together with scAAV2.7m8, which showed most efficient eGFP expression in this study. All AAV capsid variants induced eGFP fluorescence in RO and RPE after 7 days of culture (**Fig 5a**). Comparison of mean intensities of eGFP signals revealed that scAAV2.NN induced the highest expression significantly outperforming scAAV2.7m8 (**Fig 5b**). scAAV2.GL showed a comparable expression strength to scAAV2.7m8 both at day 3 and day 7. The comparison of the non-organoid, RPE area of the respective chips rendered similar results: Again, the eGFP signal of scAAV2.NN transfected RoC wells was significantly higher than the other conditions at both day 3 and 7.

**Figure 5.**
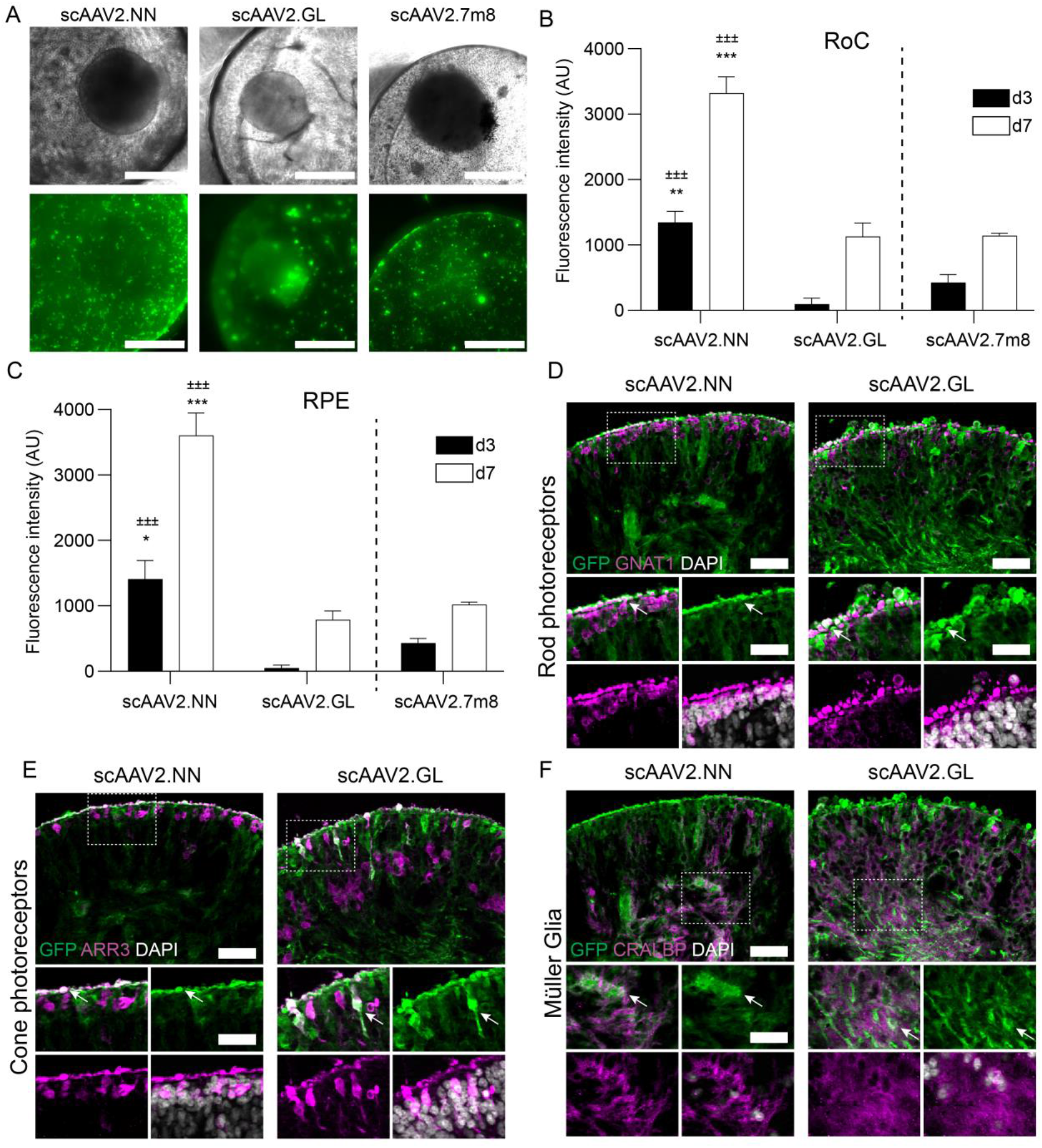
Evaluation of next generation AAVs in the RoC.(A) Representative brightfield and eGFP fluorescence (maximum intensity projections) images of RoCs (1E+10 vector genomes per well). (B) Quantification of the mean eGFP fluorescence. AAV2.7m8 data (**Fig 3-4**) is depicted as comparison. Statistical analysis represents the comparison with scAAV2.GL (±) and scAAV2.7m8 (*). n=3 for all conditions, Mean + SEM. (C) Quantification of the mean eGFP fluorescence in the non-organoid area of the RoC well. Statistical analysis represents the comparison with scAAV2.GL (±) and scAAV2.7m8 (*). n=3 for all conditions, Mean + SEM. (D-F) Vertical cryo-sections of day 300 ROs transduced with 1E+10 virus genomes showing eGFP expression (green) and cellular co-stainings (magenta). Hoechst: white. Co-stained cells are highlighted with white arrow. Scale bars: 50 µm (large images), 30 µm (small images). See also **Figure S6**.

Assessing the cellular transduction profile of the 2^nd^ generation AAVs in the RoC with day 300 ROs, we found that both AAVs were able to transduce rod and cone photoreceptors, Müller glia and bipolar cells (**Fig 5d-f, Fig S7b**). In contrast to that, none of the tested serotypes induced GFP expression in horizontal cells (**Fig S7a**). Amacrine cells and ganglion cells were not detected in day 300 RO and therefore not assessed.

### The RoC enables assessment of long-term effects of AAV transduction

One major advantage of the RoC is its long-term stability that allows analysis of the AAV induced expression over a prolonged period of time. This is of particular importance to mimic development after application, kinetics of viral transcription but also to gain pharmacokinetic information about vector-based delivery of therapeutics. In the RoC, AAV vectors can be added to the upper well which can be sealed after application, thus creating an enclosed stable system nurtured by the medium provided by the vasculature-mimicking bottom channel (cf. **Fig 3a**). To demonstrate the ability of long-time monitoring, we cultured the RoC for 21 days in presence of 3 different viral vectors (ssAAV8, ssAAV2 and ssAAV2.7m8, **Fig 6**) and monitored eGFP signals after 3, 7, 14 and 21 days. SsAAV2 and ssAAV2.7m8 showed only a faint fluorescence signal at day 7, which only slightly increased until day 21 (**Fig 6a**). This is in line with ss vectors being less potent and having a lower onset kinetic then sc vectors. At the same time, ssAAV8 eGFP showed a strong eGFP signal on day 7 continually increasing over the course of 21 days, resulting in the highest expression of the 3 tested AAVs. When plotting the eGFP expression against the maximum expression at day 21, all 3 AAV vectors were associated with a steep increase of eGFP signals until day 21. Interestingly, eGFP signal of scAAV2.7m8 increased especially strong between day 3 and 7 (**Fig 6b**). In brief, we could demonstrate that the RoC can be used to continually monitor transgene production in human retinal cells for an extended period of time.

**Figure 6.**
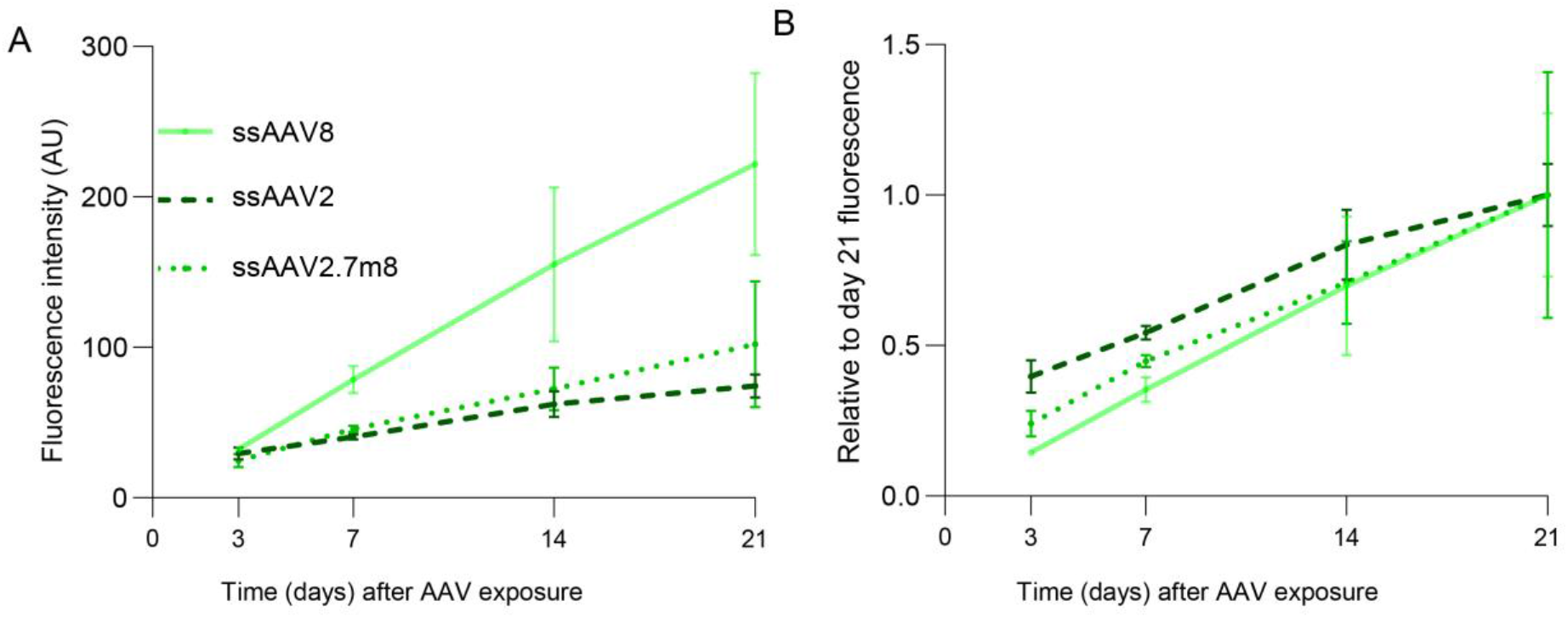
Long-term analysis of AAV induced expression in the RoC using ssAAV8, ssAAV2 and ssAAV2.7m8.(A) Quantification of the mean eGFP fluorescence in the RoC with day 300 ROs exposed to 1E+10 virus genomes per well. n=3 for all conditions, Mean ± SEM. (B) Data from (A) depicted as proportion of day 21 fluorescence level.

## Discussion

The advent of human stem cell derived microphysiological systems paralleled by AAV based gene therapies maturing into actual therapies led to the obvious and appealing concept of combining these two fast evolving fields to foster the development of novel AAV vectors and efficient gene therapies. Being immune privileged, highly compartmentalized, with good accessibility and low number of cells to be targeted, the eye is the ideal organ for a gene therapeutic approach. Hence, retinal diseases are a main focus of AAV-based research for which the versatility, availability and relevance of human ROs hold great value.

Previous reports indicated the feasibility and validity of this approach, comparing AAV transduction efficacy on stem cell-derived RPE and photoreceptors (Gonzalez-Cordero et al., 2018) and analyzing mechanisms of cell attachment and their interplay with primary cell-surface receptors (Hernandez et al.2020). AAVs were even successfully used to correct congenital retinopathy phenotypes in patient stem cell-derived ROs (Kruczek et al., 2021). Here, we further extended this concept by an in-depth analysis using two vectors AAV2 and AAV2.7m8 which are currently applied in clinical retinal therapies as well as newly developed variants with reported tropism or increased efficacy in preclinical models (Klimczak et al., 2009; Pavlou et al., 2021). To recapitulate the complex tissue structure of the human retina and physiological drug delivery routes better than what can be achieved with ROs, we employed a RoC model. The RoC provides several advantages including a physiological interaction of the RO and RPE, compartmentalization and vasculature-like perfusion. The latter aspect not only enables a constant nutrient supply mimicking choroidal vasculature but also provides the ability to deliver compounds in a subretinal-like manner. Thus, the AAV particles can be added to the upper well and retained by the epithelial barrier. In addition, the optical accessibility and stability of the RoC provides the opportunity for extended *in situ* live-cell imaging.

In the initial step of this study, we confirmed the efficacy of our AAV vector material in an animal model *in vivo*. Therefore, we applied the well described serotypes; ssAAV2, harboring a single stranded sequence of eGFP, as well as scAAV2, scAAV2.7m8, and scShH10, harboring a self-complementary sequence of eGFP, all under the control of the ubiquitous promotor CMV intravitreally to adult mice. Obtained data confirmed the expected increased expression of scAAV2s compared to ssAAV2 (McCarty et al., 2001) as well as the superior transduction efficacy of scAAV2.7m8 compared to scAAV2 (Khabou et al., 2016). The published preferential tropism for Müller glia in rodents of the ShH10 variant, a vector generated from an AAV6 parent serotype shuffled (ShH) library (Klimczak et al., 2009), could also be reproduced and was especially prominent at lower vector doses, demonstrating full functionality of the produced AAV vectors.

Second, we transferred these vectors together with the AAV9 vector to iPSC-ROs of different ages in order to assess the impact of maturation on the AAV transduction efficacy. AAV9 was added as it exhibited potent transduction after subretinal injections to rodents (Allocca et al., 2007) yet was unsuccessful to transduce retinal cells after intravitreal injections *in vivo* (Lee et al., 2019). The application of AAV9 to human retinal explant cultures from the subretinal side showed a rather low transduction efficacy (Wiley et al., 2018). In our study, scAAV9 eGFP signals were barely above the detection limit both in day 80 and day 300 ROs up to 7 days post infection. In comparison, ROs transduced with scAAV2, scAAV2.7m8 and scShH10 displayed augmented eGFP signal of which scAAV2.7m8 promoted the strongest response. Albeit using different protocols for iPSC-RO generation and AAV application, previous studies also reported poor efficacy of scAAV9 (Gonzalez-Cordero et al., 2018) and higher potency of AAV2.7m8 vs AAV2 vector (Garita-Hernandez et al., 2020). The authors of these studies indicated the lack of efficacy due to the developmental stage of their ROs (13 weeks) with missing outer segments. However, we tested AAV9 on ROs with the age of up to 42 weeks with formed inner and outer segments, suggesting that other mechanisms might be responsible for the lack of transduction.

After validating the effects of the AAV vectors on hiPSC-ROs, the same vector panel was applied to the RoC. In the RoC, scAAV2 transduced retinal cells as well as the RPE. This ability is in line with the capsid properties shown in rodent models (Auricchio et al., 2002) and essential for the therapeutic benefit of Luxturna in patients. Aside from AAV2, a more significant eGFP expression in RPE and retinal cells was induced by the scAAV2.7m8 and scShH10 vector variants. Remarkably, the reported cellular tropism of ShH10 in rodents towards Müller glia cells did not translate to the RoC. On the contrary, in the human RoC setting, the ShH10 vector shows a high potency to transduce the RO and RPE with comparable potency to the AAV2.7m8. This is in line with results from Codero et al. testing ShH10 using human iPSC-ROs. We also detected the extension of ShH10 transduction to ganglion and horizontal cells which had not been analyzed in previous studies. Notably, this non-selective tropism was also observed for the lower virus doses tested (e.g. 1E+8 vector genomes per RoC well, data not shown). Collectively these data show that the tropism found in rodents does not fully translate to models based on iPSC-derived human cells. This raises concerns whether the reported Müller glia cell tropism can be translated to the human retina. Additional data from this vector in higher species are unfortunately lacking but might give a more complete picture of this vector’s capacities.

The highest measured signals were obtained from RoC treated with scAAV2.7m8. The AAV2.7m8 capsid variant was selected from an AAV2 based peptide insertion library and has a higher potency to transduce murine and NHP retinae (Dalkara et al., 2013). Therefore, this variant was utilized as the clinical phase II candidate vector for intravitreal injection aiming for the treatment of patients with wet age-related macular degeneration and diabetic macular edema (Grishanin et al., 2019) currently in clinical phase II. In unison with the convincing data compiled for the AAV2.7m8 vector variant, our RoC model delivers supportive data for the capabilities of this newly generated AAV vector, particularly to efficiently transduce the main retinal cell types including photoreceptors, Müller glia and ganglion cells.

In addition, we interrogated whether and how newly-developed AAV vectors selected for retinal application would perform in the RoC. Therefore, we applied the two recently published variants AAV2.NN and AAV2.GL (Pavlou et al., 2021). These AAV2 based heptamer insertion mutants were reported as potent transducers of retina in mice, dogs, NHPs and human explants (Pavlou et al., 2021). To assess their performance in the RoC, AAV2.NN and AAV2.GL were compared to scAAV2.7m8 which served as an intra-experimental control. While scAAV2.GL showed comparable expression to scAAV2.7m8, scAAV2.NN significantly surpassed both, showcasing how the RoC potential can be leveraged in order to screen and detect increased potency of novel AAV variants for transduction of human retinal tissue.

Taken together, our study reveals for the first time the potential of a complex microphysiological system as a translational model for testing of AAV vectors in a clinical-like setup. We could show that the RoC offers the ability to directly compare the transduction efficacy, the cellular tropisms using a subretinal-like application route which is one of the two majorly used application routes in the clinical context. An important future implementation of the RoC will be to mimic the intravitreal application currently limited due to the spheroidal nature of the RO and the absence of an accessible inner limiting membrane. Overall, the RoC can be an important asset in the development and safety assessment of new vector candidates for future retinal gene therapies.

## Experimental Procedures

### AAV vectors

AAV2, AAV8, AAV9, AAV2.7m8 (Dalkara et al., 2013), ShH10 (Klimczak et al., 2009), and AAV2.NN, AAV2.GL (Pavlou et al., 2021) with the single stranded (ss) expression cassettes; ss-CMV-eGFP-SV40polyA, ss-CMV-antiFITC or self complementary (sc) sc-CMV-eGFP-SV40polyA, (Strobel et al., 2020) flanked by AAV2-derived inverted terminal repeats were applied. AAV vectors were prepared as previously described (Strobel et al., 2019). Briefly, HEK 293-H cells, (Thermo Fisher Scientific) were cultured in DMEM+ GlutaMAX-I + 10 % fetal calf serum (Thermo Fisher Scientific) and transfected as previously described (Strobel et al., 2015). AAV purification via polyethylenglycol precipitation, iodixanol gradient, ultrafiltration and sterile filtration was conducted as previously described (Strobel et al., 2019).

### Virus titer measurement via digital droplet PCR

AAV genomic titers were determined by digital droplet PCR using QX200 system (Bio-Rad, USA) with the following primers specific for the CMV promotor; forward: CCAAGTACGCCCCCTATTGAC. reverse: CTGCCAAGTAGGAAAGTCCCATAAG. Viral DNA was prepared with ViralXpress (Merck Millipore, USA). Droplets were generated using the Bio-Rad Droplet Generator (Bio-Rad, USA). X50s PCR Mastercycler (Eppendorf, Germany). was used with initial denaturation step 10 min at 95°C, followed by 40 cycles of 30 sec at 95°C with annealing for 1 min at 60°C. AAV genomic titers were analyzed with Droplet Reader using the QuantaSoft software (Bio-Rad, USA).

### Animal experiments

9-12 weeks old C57BL/6J female mice (Charles River, Germany) were used in this study. Experiments were approved by local animal welfare authority (Regierungspräsidium Tübingen, Germany) and conducted according to the German Welfare Law and the GV-SOLAS guidelines (Dülsner et al., 2017). Mice were anesthetized with 3.5 % isoflurane and 1 µl of AAV suspension was injected transsclerally into the vitreous using a 34G canula (WPI, USA). Animals were sacrificed under anesthesia by cervical dislocation 3 weeks after injection and eyes were harvested in 4 % paraformaldehyd for Immunohistochemistry or liquid nitrogen for quantitative reverse transcriptase-PCR.

### Fabrication of the RoC

Design and fabrication of the RoC was conducted as described previously (Achberger et al., 2019). In comparison to the previously described version, the design of the top layer was slightly modified to integrate tissue compartments with an increased volume of 27 µl.

### Cell culture

The iPSC line used in this study was derived from hair keratinocytes of a healthy male donor as previously described (Linta et al., 2012). iPSCs were cultured on Matrigel (hESC-qualified, BD Biosciences, USA)-coated plates with FTDA medium (Frank et al., 2012). Cells were passaged every 6–7 days using Dispase (Stemcell Technologies, Canada). Differentiated colonies were removed manually by scraping. All procedures were in accordance with the Helsinki convention and approved by the Ethical Committee of the Eberhard Karls University Tübingen (Nr. 678/2017BO2). The control person gave his written consent.

HiPSC-ROs were differentiated and cultured based on a protocol by Zhong et al. (Zhong et al., 2014) with some modifications as previously described (Achberger et al., 2019). RPE cells were derived as a product from RO differentiation following (slightly adapted) procedures of Zhong et al. (Zhong et al., 2014) and Ohlemacher et al. (Ohlemacher et al., 2015) and differentiated and maintained as previously reported (Achberger et al., 2019). RoCs were assembled and cultured as previously described (Achberger et al., 2019). In contrast to the previous reported protocol, RPE were cultured for 2 weeks prior to the loading of ROs with daily medium change.

### AAV treatment

#### RO

One RO per well was placed in non-adherent 48 well plates in 300 µl BRDM on day -1. On day 0, AAVs were thawed and the required amount of AAV was added by a 33,3 % medium change. After 1, 2 and 3 days of AAV exposure, a 100 % medium change was performed including washing with Dulbecco’s phosphate-buffered saline (PBS, no calcium, no magnesium, Thermo Fisher Scientific, USA). An additional medium change was done on day 5.

#### RoC

The AAV treatment consisted upon replacing the volume of medium with the AAV solution/mix containing the desired virus genome number. The volume added was between 1-8 µl. The final volume of the tissue compartment was kept to 27 µl, defined by the chip design.

### Immunohistochemistry

Whole eyes were fixed in 4 % PFA and paraffin embedded (formalin fixed and paraffin embedded, FFPE). 3 µm thick sections of FFPE tissue on super frost plus samples were deparaffinized and rehydrated by serial passage through changes of xylene and graded ethanol for immunohistochemistry staining. Antigen retrieval was performed by incubating the sections in Leica Bond Enzyme solution (Bond Enzyme Pre-treatment Kit, Leica Biosystems, Germany) for 5 min. Sections were incubated (30 min, room temperature (rt)) with an anti-GFP antibody (see **Table S2**) in Leica Primary Antibody Diluent (Leica Biosystems). Bond Polymer Refine Detection (Leica Biosystems) was used for detection (3,3‘Diaminobenzidine as chromogen, DAB) and counterstaining (hematoxylin). Staining was performed on the automated Leica IHC Bond-III platform (Leica Biosystems).

The RoC was disconnected from the perfusion system and washed with PBS through the medium channel and in each well. Fixation was performed with 4% Histofix (Carl Roth, Germany) for 1 hour at rt and with mild agitation. After fixation, the RoC were washed with PBS and kept at 4°C until embedding. 1 day before embedding, RO were retrieved from RoC by flushing the chip wells with PBS. The collected RO were kept in 30 % Sucrose (in PBS) over night, embedded in cryomolds using Tissue-Tek O.C.T. (Sakura Finetek, USA) and frozen in -80°C until cryosectioning. Cryosections were performed at a cryostat (14 µm slices, CM 3050 S Cryocut, Leica Biosystems), mounted on Superfrost microscope slides (Thermo Fisher Scientific, USA). Before staining, slides were rehydrated in PBS for 15 min and incubated in a blocking solution of 10 % normal donkey serum in PBS with 0.2 % triton-X for 1 hour. Primary antibodies (**Table S2**) were diluted in blocking solution and incubated overnight at 4°C. Secondary antibodies (Abcam, UK) were diluted in 1:1 blocking solution:PBS and incubated for 2 hours at rt. Mounting was performed with ProLong Gold Antifade Reagent (Thermo Fisher Scientific, USA). Washing steps to remove residual antibodies were performed with PBS, three times for 3 min after each antibody incubation. Hoechst 33342 (1:2000, Thermo Fisher Scientific, USA) was added to the last washing step.

### Gene expression analysis

For RNA isolation, samples were homogenized in 900 μL RLT buffer (Qiagen, Germany), using a Precellys 24 homogenizer and ceramic bead tubes (VWR, USA) at 6000 rpm for 30 sec. Afterwards, samples were immediately placed on ice. 350 μl Phenol-chloroform-isoamyl alcohol (Sigma Aldrich, USA) were then added to 700 μl homogenate in a phase lock gel tube and mixed by shaking. Following centrifugation for 5 min at 16000 × g, 350 μl Chloroform-isoamyl alcohol (Sigma-Aldrich, USA) were added and the mixture was shaken again. After 3 min of incubation at rt and centrifugation for 5 min at 12000 × g, the upper phase was collected and pipetted into a deep well plate placed on dry ice. After processing of all samples, DNA and RNA were purified, using the AllPrep DNA/RNA 96 kit (Qiagen, Germany) as per instructions, including the optional “on-column DNase digestion” step. Integrity of RNA was analyzed by BioAnalyzer (RIM values>6) AAV vector genomes were measured using extracted DNA and a standard curve generated by serial dilutions of the respective expression plasmid. Taqman runs were performed on an Applied Biosystems ViiA 7 Real-Time PCR System (ThermoFisher Scientific, USA). For gene expression analysis, equal amounts of RNA were reversely transcribed to cDNA using high-capacity cDNA RT kit (Applied Biosystems, ThermoFisher Scientific) as per instructions. qRT-PCR reactions were then set up using RT-PCR kit (Applied Biosystems, ThermoFisher Scientific). Primers used are listed in **Table S3**.

The RoC was disconnected from the perfusion and washed with PBS though the medium channel and on each well. The bottom layer and the membrane were removed for collection of the RPE using a scalpel. The RO within the hydrogel was collected in 500 µl of PBS. Dry samples were shock frozen in liquid nitrogen. RNA was isolated by pelleting cells, followed by lysis in 350 μl RLT buffer and purification using the RNeasy mini kit (74104, Qiagen) according to manufacture’s instruction. Primers used are listed in **Table S3**.

### Microscopy

Microscopic assessment of mice sections was conducted with a Zeiss AxioImager M2 microscope and ZEN slidescan software (Carl Zeiss, Oberkochen, Germany). Plate-cultured ROs were imaged with a Zeiss Axio Imager.Z1 (Carl Zeiss). RoCs were imaged with a Leica DMi8, LEICA Microsystems GmbH, Germany. Cryosectioned RO extracted from RoC were imaged using an Imager.M2 Apotome1 (Carl Zeiss). RoC confocal images were shot using a Zeiss LSM 710 version 34ch Quasar NLO equipped with the inverted microscope platform Axio Observer.Z1.

### Image analysis

Fluorescence quantification of plate-cultured ROs was conducted using a macro (supplementary macro 1) in ImageJ (imagej.nih.gov). Briefly, the macro determined the RO-covered area using a threshold on the bright field image and the Plugin “Analyze Particles”. The mean grey values of the selected area was then analyzed in the respective eGFP channel using the Measure plugin of ImageJ.

For fluorescence quantification in the RoC, ImageJ (Supplementary macro 2) was used to generate the projection (sum) for both brightfield and GFP channels. The area of each RO was selected manually in the brightfield image. The mean grey values were quantified within this area using the Measure function of ImageJ. All fluorescence values were adjusted for background. For both ROs and RoC, the selection areas allowed to monitor the average diameters and evaluate possible morphological changes.

### Statistics

Statistical analysis was performed with GraphPad Prism 9.0 (Graphpad Software, USA) and statistical testing was performed using one-way ANOVA with Bonferroni post-hoc test (**Fig 1** and **Fig 3c**) and two-way ANOVA with Bonferroni post-hoc test (**Fig 1a, Fig 2b, Fig 3c, Fig 3d, Fig 5b, Fig 5c**). *p<0.05, **p<0.01, ***p<0.001.

## Acknowledgements

Cristhian Rojas (Fraunhofer IGB) for chip production, Kirstin Linke (IGB) for cell seeding on the chip, Anamaria Bernal Vergara (EKUT) for histological sample preparation and Laura Laistner (IGB) for image acquisition.

## Author Contributions

Conceptualization, P.L., M.D., C.S., St.K., Se.K., M.C., T.L., S.M., U.M., A.K., S.L. and K.A.; Methodology, M.D., C.S., M.C., K.A., A.K. und J.C; Investigation M.C., M.D., J.R, L.M., V.C., S.P, N.P, B.S., and S.C; Writing K.A, M.D., P.L., M.C., A.K. and C.S.; Resources, T.L, U.M., P.L und S.L.; Supervision, P.L., S.L., Se.K, M.D., and U.M.

