## Supplementary Material for "Human stem cell-based retina-on-chip as new translational model for validation of AAV retinal gene therapy vectors"

**Supplemental figures**

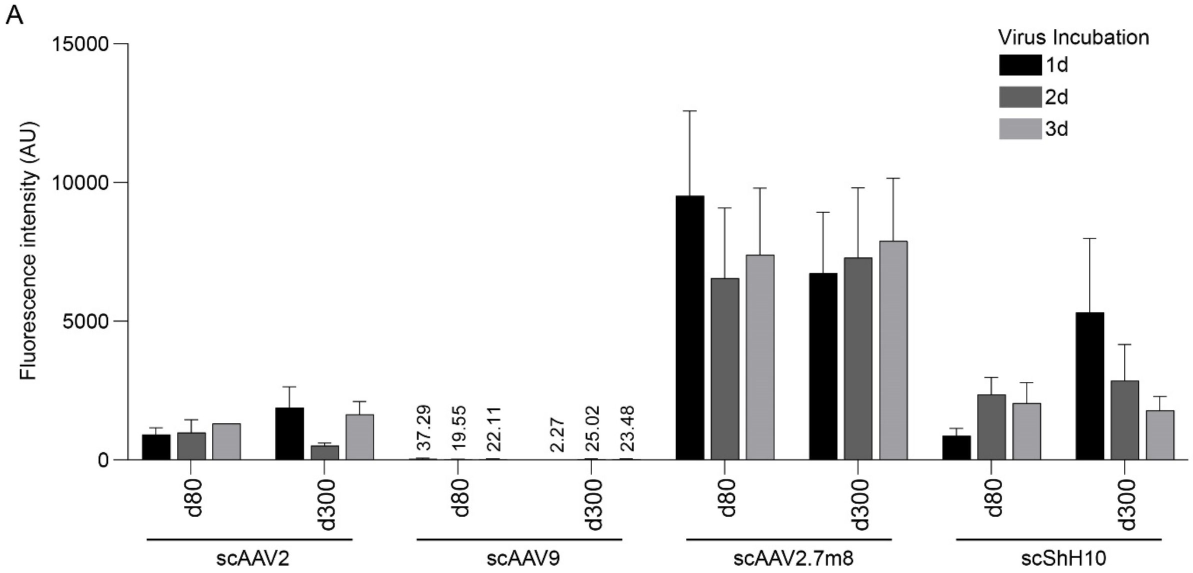


**Figure S1.** eGFP expression on AAV treated retinal organoids differentiated for 80 and 300 days separated by inoculation time, **Related to Figure 2**.

(A) Quantification of the mean eGFP fluorescence in day 80 and day 300 retinal organoids exposed for 1, 2 or 3 days to the respective AAV (1E+10 virus genomes/well) and quantified after 7 days. Numbers above the bars give the mean eGFP fluorescence for the respective conditions. There is no statistical difference between the 3 incubation times for all conditions (Two-Way ANOVA with Bonferroni post-hoc test). n=5-6 per condition, Mean + SEM.

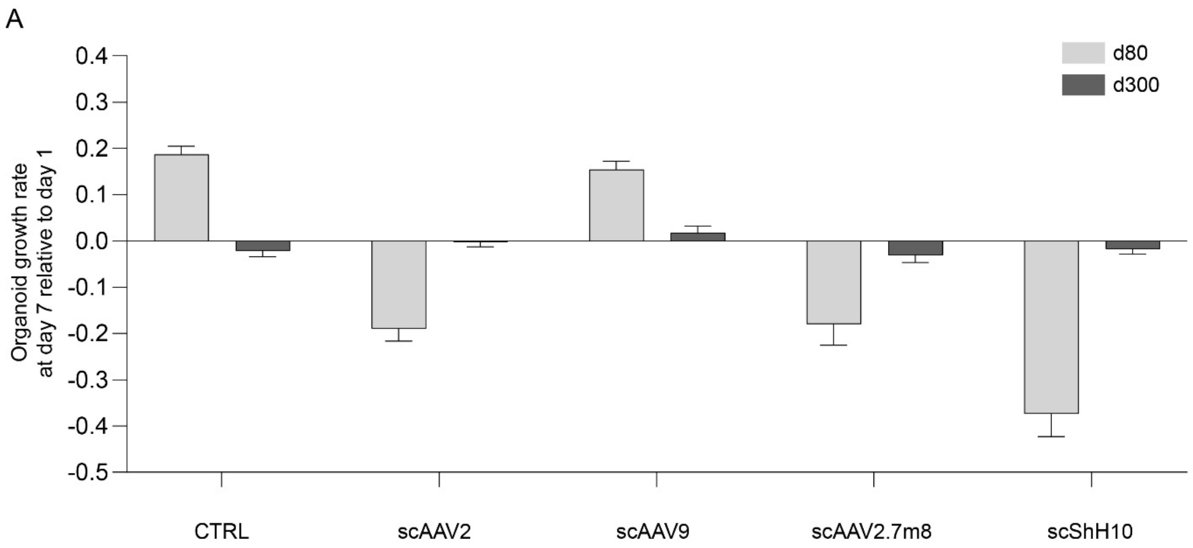


**Figure S2**. Relative size change of retinal organoid during the AVV transduction, **Related to Figure 2**.

(A) Graphs shows relative retinal organoid size comparing day 7 and day 1 after AAV transduction. Controls were untransfected. n=6 for control and n=18 for AAV treated organoids, Mean + SEM.

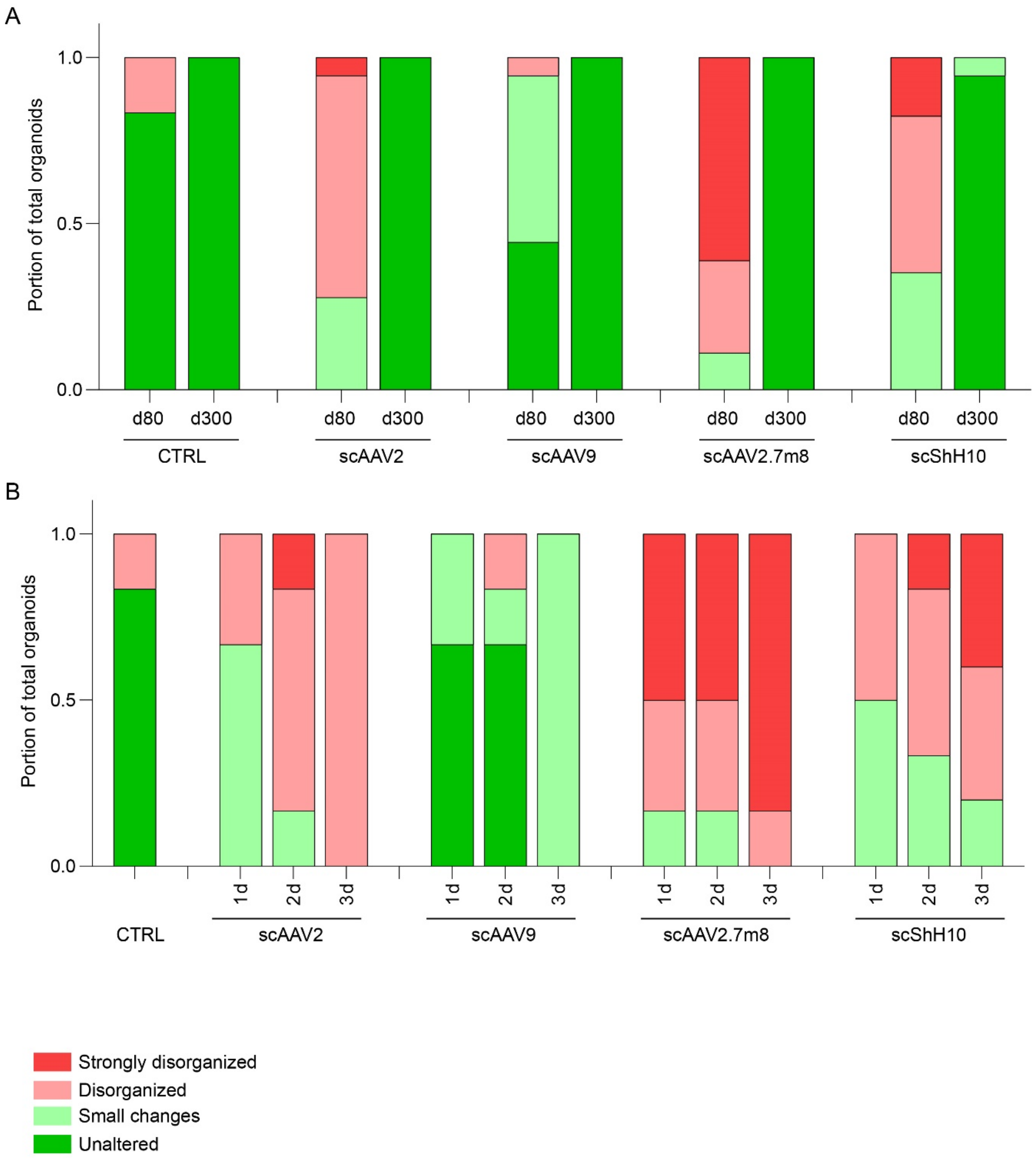


**Figure S3**. Integrity assessment of retinal organoids after 7 days of AAV transduction, **Related to Figure 2**.

(A) Morphological assessment of day 80 and day 300 retinal organoids for non-treated control (CTRL) and transduced with the AAV serotypes. Criteria were as indicated in the legend. Graph shows the proportion of the total assessed retinal organoids for each morphological score. n=6 for controls and 17-18 for treated conditions.

(B) Organoid morphological of day 80 retinal organoids discriminated by inoculation time (1 day, 2 days and 3 days). n=5-6 for all conditions.

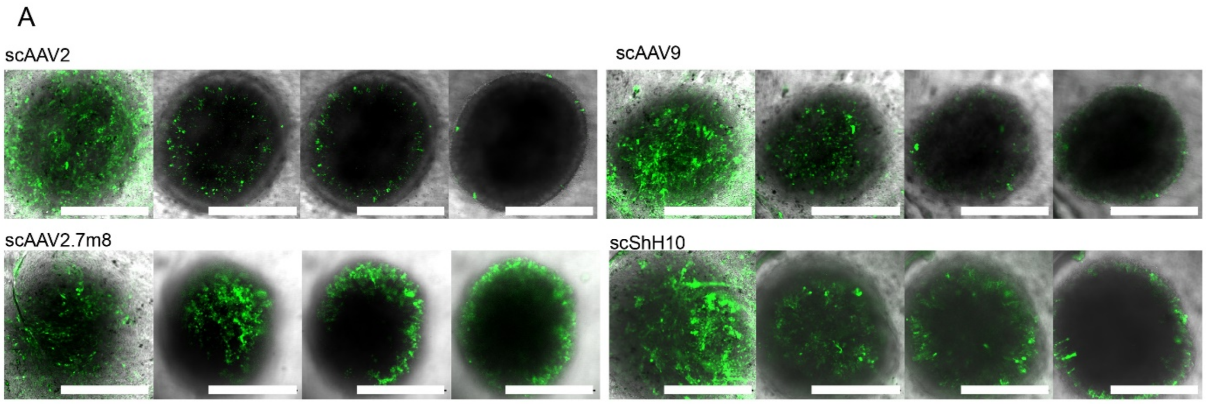


**Figure S4**. Confocal imaging of AAV-transduced RoC. **Related to Figure 3**.

(A) Representative brightfield and confocal eGFP fluorescence live imaging on RoC with day 80 and 300 retinal organoids and day 300. The fluorescent images are presented as maximum intensity projections. Scale bars: 500 μm.

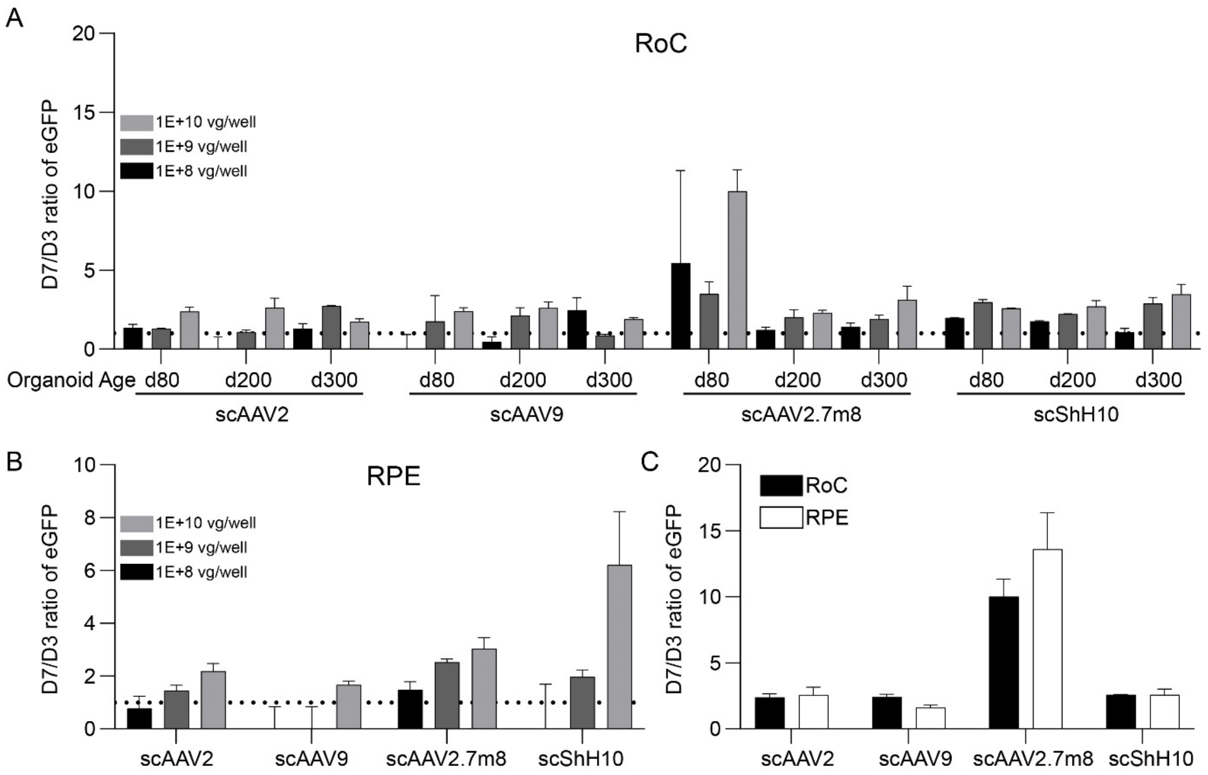


**Figure S5.** Kinetics of AAV transduction in the retina-chip (RoC), **Related to Figure 3.**

(A) Relative eGFP signal comparing day 3 and day 7 after transduction with the respective AAV in the RoC discriminated by the retinal organoid age (d80, d200 and d300). Virus load as indicated in the figure legend. n=3 per condition, mean + SEM. (B) Relative eGFP signal comparing day 3 and day 7 after transduction with the respective AAV in the non-organoid area of the RoC wells. Virus load as indicated in the figure legend. n=3 per condition, Mean + SEM. (C) Relative eGFP signal comparing day 3 and day 7 after transduction with the respective AAV in the RoC and in the non-organoid area of the RoC wells (RPE) treated with the indicated AAV, with a virus load of 1E+10 virus genomes per well. n=3 for RoC and n=5 for RPE, Mean + SEM.

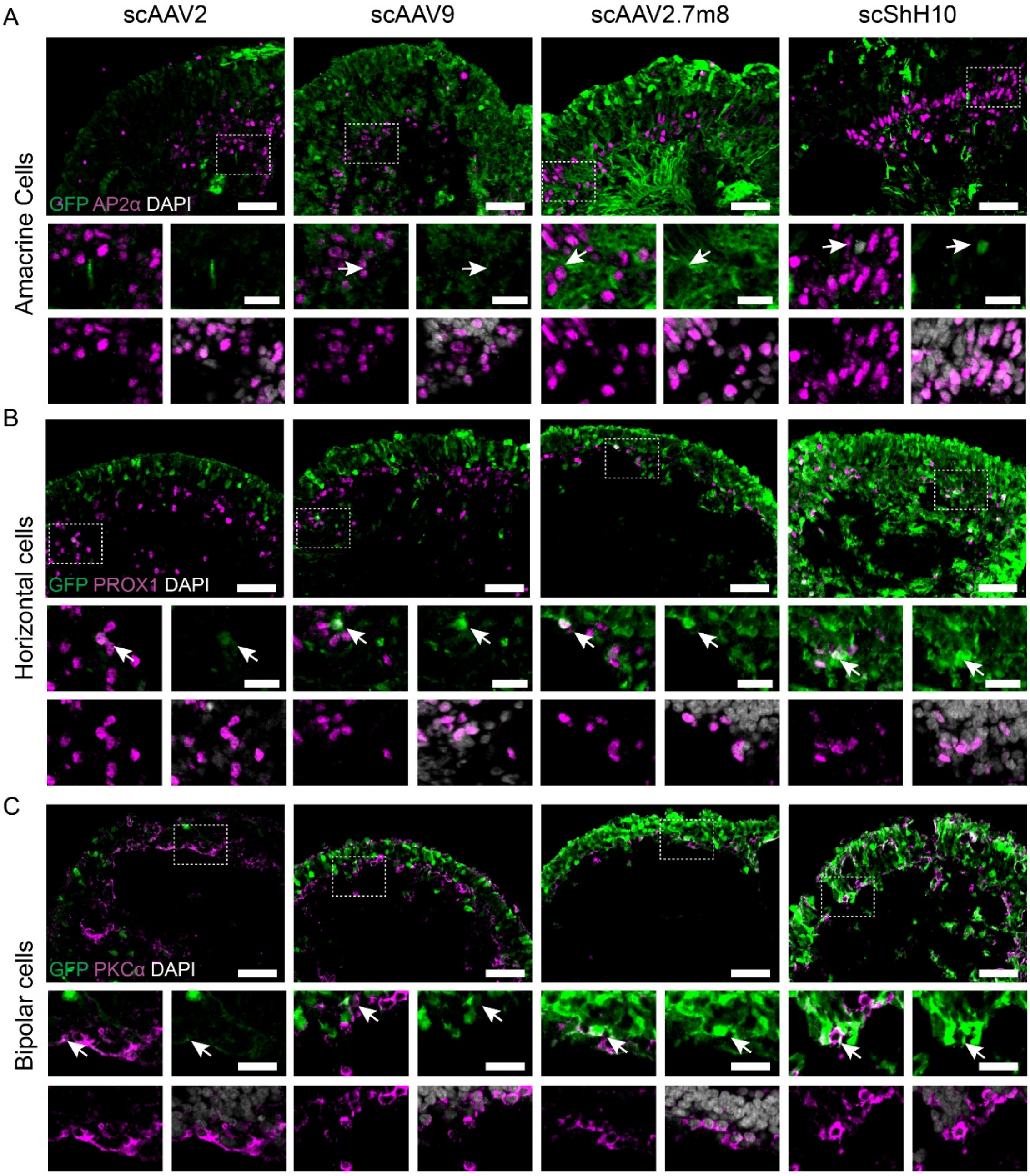


**Figure S6**. Additional data of cell tropism of AAV2, AAV9, AAV2.7m8 and ShH10 in the retina-chip (RoC), **Related to Figure 3**.

Retinal organoids in (A) at day 80 were transduced with 1E+10 virus genomes, day 200 (B-C) retinal organoids were transduced with 1E+10 virus genomes. (A-D) Vertical cryo-sections of organoids showing AAV-mediated eGFP expression (green) and cellular co-stainings (magenta) for AP2α (A, Amacrine Cells), PROX1 (B, Horizontal cells) and PKCα (C, Bipolar Cells). Cell nuclei were stained with DAPI (white). Co-stained cells are highlighted with white arrow. Dotted squares indicate the position of the 4 magnified areas shown below each image. Scale bars: 50 µm (large images), 20 µm (small images).

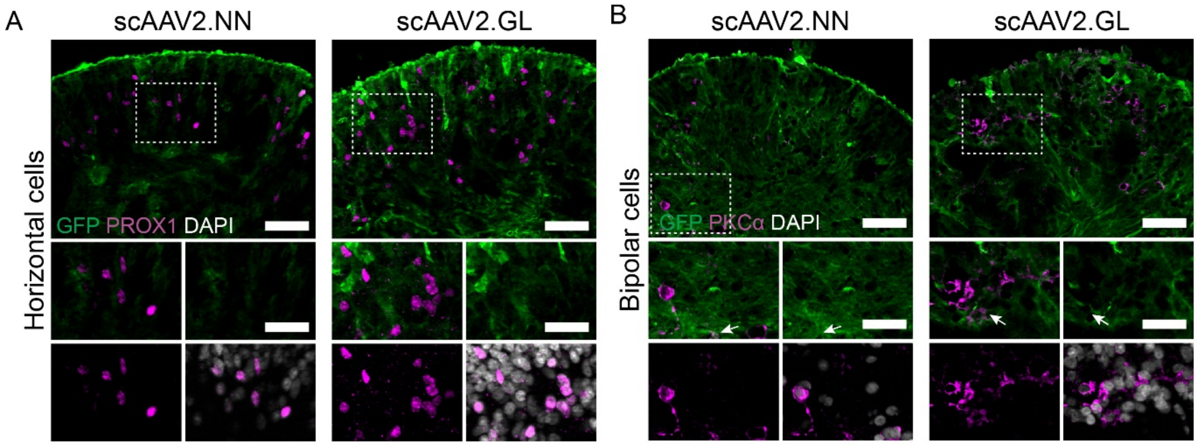


**Figure S7**. Evaluation of cell tropism in horizontal and bipolar cells of new generation AAVs in the RoC, **Related to Figure 5**.

(A-B) Vertical cryo-sections of day 300 retinal organoids transduced with 1E+10 virus genomes showing AAV2.NN and AAV2.GL mediated eGFP expression (green) and cellular co-stainings (magenta) for PROX1 (A, Horizontal Cells) and PKCα (B, Bipolar Cells). Cell nuclei were stained with DAPI (white). Co-stained cells are highlighted with white arrow. Dotted squares indicate the position of the 4 magnified areas shown below each image. Scale bars: 50 µm (large images), 30 µm (small images).

**Supplemental tables**

**Table S1**. Summarized results of all tested AAV serotype.

| | | scAAV2 | scAAV9 | scAAV2.7m8 | scShH10 | scAAV2.NN | scAAV2.GL |
|---|---|---|---|---|---|---|---|
| GFP expression | RNA RO | + | + | +++++ | +++ | NA | NA |
| | Fluorescence ROC | ++ | + | ++++ | +++ | +++++ | ++++ |
| | Fluorescence RPE | ++ | + | ++++ | +++ | +++++ | ++++ |
| Cell Tropsim | Cones | (green) | (green) | (green) | (green) | (green) | (green) |
| | Rods | (green) | (green) | (green) | (green) | (green) | (green) |
| | Müller Glia | (green) | (green) | (green) | (green) | (green) | (green) |
| | Ganglion Cells | (green) | (green) | (green) | (green) | NA | NA |
| | Amacrine Cells | (yellow) | (green) | (green) | (yellow) | NA | NA |
| | Bipolar Cells | (red) | (red) | (green) | (green) | (yellow) | (yellow) |
| | Horizontal Cells | (green) | (green) | (green) | (green) | (red) | (red) |

- (green) Several positive cells
- (yellow) Only 1-2 positive cells
- (red) No positive cells

NA: not available

**Table S2**. List of primary antibodies used

| Antibody | Dilution | Catalogue number | Company |
|---|---|---|---|
| GFP (whole eye paraffin sections) | 1:1500 | Ab290 | Abcam, UK |
| GFP (RO sections | 1:1500 | GFP-1010 | Aves Labs, USA |
| AP2α | 1:100 | Sc-12726 | Santa Cruz Biotechnology, USA |
| Arrestin 3 | 1:50 | Sc-54355 | Santa Cruz Biotechnology, USA |
| BRN3B | 1:50 | Sc-31989 | Santa Cruz Biotechnology, USA |
| CRALBP | 1:250 | Ab15051 | Abcam, UK |
| PKCα | 1:500 | Sc-208 | Santa Cruz Biotechnology, USA |
| GNAT1 | 1:500 | GTX114440 | GeneTex, USA |
| PROX1 | 1:2000 | ABN278 | Merck Millipore, USA |

**Table S3**. List of genes, sequences of primers and Taqman probes used for RT-qPCR

| Gene | Forward | Reverse | Probe |
|---|---|---|---|
| eGFP | CTGCTGCCCGACAACCA | TGTGATCGCGCTTCTCGTT | TACCTGAGCACCCAGTCCGCCCT |
| Murine RNA polymerase II (*Polr2a*) | GCCAAAGACTCCTTCACTCACTGT | TTCCAAGCGGCAAAGAATGT | TGGCTCTTTCAGCATCTCGTGCAGATT |
| human RNA polymerase II (*RPB1*) | GCAAGCGGATTCCATTTGG | TCTCAGGCCCGTAGTCATCCT | AAGCACCGGACTCTGCCTCACTTCATC |
